# Linker Histone H1 subtypes specifically regulate neutrophil differentiation

**DOI:** 10.1101/763482

**Authors:** Gabriel Sollberger, Robert Streeck, Brian E. Caffrey, Arthur I. Skoultchi, Arturo Zychlinsky

## Abstract

Neutrophils are important innate immune cells that tackle invading pathogens with different effector mechanisms. They acquire this antimicrobial potential during their maturation in the bone marrow, where they differentiate from hematopoietic stem cells in a process called granulopoiesis. Mature neutrophils are terminally differentiated and short-lived with a high turnover rate. Here, we show a critical role for linker histone H1 on the differentiation and function of neutrophils using a genome-wide CRISPR/Cas9 screen in the human cell line PLB-985. We systematically disrupted expression of somatic H1 subtypes to show that individual H1 subtypes affect PLB-985 maturation in opposite ways. Importantly, H1 subtypes also affect neutrophil differentiation of murine bone marrow stem cells, demonstrating an unexpected subtype-specific role for H1 in granulopoiesis.

## Introduction

Because multicellular organisms often face disturbances in their homeostasis, they evolved complex immune responses to deal with sterile and infectious insults. Like all cells in the human body, immune cells differentiate from multipotent precursors, which in the immune context are primarily hematopoietic stem cells in the bone marrow. One of these differentiation programs, called granulopoiesis, leads to the development of granulocytes - basophils, eosinophils and neutrophils. Neutrophils are the most abundant leukocytes in humans. In homeostatic conditions, up to 2×10^11^ neutrophils enter the blood stream per day and patrol the host’s body until they sense signs of infection, which triggers them to leave the blood stream and migrate to the inflammatory site where they ensure pathogen removal. An efficient neutrophil response is crucial for human antimicrobial defense and, correspondingly, neutropenia is associated with severe infections^1^.

The three best described defense mechanisms of neutrophils are phagocytosis (engulfment of pathogenic microorganisms and subsequent destruction in phagosomes), degranulation (the release of antimicrobial proteins from granules to the extracellular space) and the formation of neutrophil extracellular traps (NETs). NETs are chromatin structures studded with antimicrobial proteins derived mostly from neutrophil granules, which can trap pathogens^2^. Neutrophils release NETs via various mechanisms, which in most cases lead to neutrophil death^3^; thus, NETs are an extreme form of antimicrobial defense, as their release also eliminates the cell casting the NET. NET formation occurs in mature neutrophils and therefore we propose that specific cues during neutrophil differentiation shape their ability to form NETs.

Neutrophils differentiate from granulocyte-macrophage precursors. The first progenitor with a clear neutrophil lineage preference during granulopoiesis is the promyelocyte^4^. During maturation, neutrophils acquire a lobulated nuclear shape and produce many of their antimicrobial effector proteins, stored in granules^5^. Granulopoiesis requires balanced activity of various transcription factors^4^. The C/EBP family, especially C/EBP-α and –ε, are key regulators of neutrophil differentiation^6, 7^. Other transcription factors can antagonize C/EBP-dependent neutrophil development. For example, expression of GATA-1^8^ and GATA-2^9^ induce eosinophil, basophil or mast cell differentiation^10^.

Differentiation programs rely both on transcription factor activation and on epigenetic chromatin alterations. DNA wraps around an octamer of core histones forming a nucleosome. The linker histone H1 binds to nucleosomes and further compacts chromatin^11^. There are 11 different H1 subtypes in mice and humans. Five of them (H1.1 – H1.5) are somatic replication-dependent subtypes, two (H1.0 and H1X) are somatic and replication-independent and four (H1oo, H1t, H1T2 and HILS1) are germline-specific^12^. All H1 subtypes consist of a highly conserved globular domain and two more variable regions: the N- and the C-terminal tails. Individual H1 subtypes are more conserved between species than H1 subtypes are within one species, suggesting that the conservation of different subtypes has functional relevance^12^. Still, some functions of H1, such as global shaping of chromatin structure, are redundant between subtypes. This is reflected by the fact that mice deficient for one or even two H1 subtypes are viable and fertile without striking morphological abnormalities^13^. However, mice lacking three H1 subtypes die *in utero*, with total H1 levels reduced to about 50%^13^. Appropriate H1 expression is therefore essential for development. Despite some redundancy, H1 subtypes have specific functions. H1 subtypes bind to chromatin with different affinities and knock down of specific H1 subtypes affects gene transcription in distinct ways^14–16^.

The role of H1 and H1 subtypes in immunity is poorly understood. H1 contributes to the silencing of pro-inflammatory cytokines after endotoxin challenge^17^, as well as to the silencing of interferon-stimulated genes^18^. Interestingly, mice deficient in H1.0 have fewer dendritic cells, but normal numbers of granulocytes, macrophages and lymphocytes, showing a subtype-specific role for H1 in dendritic cell differentiation^19^. However, to date, no studies have systematically addressed H1 subtype involvement in the maturation and function of human immune cells

Here we used the human neutrophil-like cell line, PLB-985, to study neutrophil maturation and function. We performed a genome-wide CRISPR/Cas9 screen using survival to a NET-inducing stimulus, phorbol 12-myristate 13-acetate (PMA), as a readout. Surprisingly, we found that depletion of either H1.2 or H1.4 strongly reduced the differentiation of PLB-985 and thereby their ability to form NETs. Notably, deficiency of the other somatic H1 subtypes, H1.1, H1.3 and H1.5, accelerated PLB-985 maturation. RNA-seq analysis showed that H1.2 and H1.4 deficiency leads to an upregulation of eosinophil genes in PLB-985. Mirroring these results, murine bone marrow stem cells from H1.2/H1.4 double deficient mice shifted their differentiation profile from neutrophil towards eosinophil cell fate. We further demonstrate that the subtype-specific function of H1 in neutrophil and eosinophil differentiation depends – at least in part – on the transcription factor GATA-2. We uncovered that, unexpectedly, H1 subtypes affect lineage specification during granulopoiesis.

## Materials and methods

### Cell lines

PLB-985 were kindly provided by Prof. Mary Dinauer, Washington University School of Medicine^20^.

### Antibodies and staining reagents

H1.2 antibody (PA5-32009, Invitrogen), H1.4 antibody (41328S, Cell Signaling), MPO antibody (A0398, Dako), Histone H3 antibody (ab1791, Abcam), SYTOX Green nucleic acid stain (Thermo Fisher Scientific) and 4′,6-diamidino-2-phenylindole (DAPI; Sigma) were used. All flow cytometry antibodies were from BD Biosciences: human Cd11b (557321), mouse Cd11b (562605), mouse Ly-6G (551461), mouse SiglecF (565527), mouse Cd45 (561487), mouse Cd3 (561798), mouse Cd4 (552775), mouse Cd8a (561093), mouse Cd115 (565249), mouse GR1 (561103).

### Chemicals and cytokines

Luminol (11050, AAT-Bioquest), horseradish peroxidase (31941, Serva), PMA (P8139, Sigma), A23187 (Santa Cruz Biotechnology Inc.), GATA-1 inhibitor (anagrelide, SML0846, Sigma), GATA-2 inhibitor (K-7174, HY-12743A, MedChemExpress) were used. Recombinant murine cytokines (IL-3, IL-5, IL-9, GM-CSF, SCF) were from Peprotech.

### Plasmids

All CRISPR/Cas9 gene disruptions were done using lentiCRISPRv2^21, 22^. LentiCRISPRv2 was a gift from Feng Zhang (Addgene plasmid #52961; http://n2t.net/addgene:52961; RRID:Addgene_52961). Lentiviral particles were produced using psPAX and pMD2.G packaging plasmids in HEK293T cells.

### sgRNA sequences and primers

All sgRNA sequences and primer sequences are listed in Supplementary Table 3.

### Donor consent

Human primary neutrophils, PBMCs and monocytes were isolated from blood samples of healthy volunteers according to the declaration of Helsinki. All donors provided written informed consent and all blood samples were collected with approval from the local ethics committee.

### Mice

Breeding of mice and isolation of blood and bone marrow were approved by the Berlin state authority Landesamt für Gesundheit und Soziales. All mice were bred at the Max Planck Insitute for Infection Biology under specific pathogen-free conditions. Animals were maintained on a 12-hour light/12-hour dark cycle and fed ad libitum. H1.2/H1.4-deficient animals were provided by Prof. Arthur Skoultchi^13^.

### Primary cell isolation

Neutrophils were purified by centrifugation of whole blood over Histopaque-1119 (Sigma), followed by centrifugation over a discontinuous Percoll gradient^23^. PBMCs were collected from the upper phase after Histopaque-1119 centrifugation, washed and either used for RNA extraction or for isolation of monocytes by positive selection (CD14) with magnetic beads (130-050-201, Miltenyi Biotec).

### PLB-985 propagation and differentiation

PLB-985 were cultured in RPMI (RPMI 1640 medium, gibco) supplemented with 10 % FCS, L-glutamine and antibiotics (25030-024 and 15140-122, gibco), cells were passaged every 3-4 days. For differentiation, cells were counted at d0 and 0.4×10^6^ cells/ml were seeded in differentiation medium (RPMI supplemented with 2.5 % FCS, 0.5% dimethylformamide (DMF, D4551, Sigma), L-glutamine and antibiotics), usually in 6 well plates in a total volume of 3ml. At d4 of differentiation, 2ml of differentiation medium were added per 6 well. At d7, cells were washed and passed over an equal volume of Histopaque-1077 (Sigma) (centrifugation for 20min at 800 x g, brakes off) to remove debris and subsequently seeded at the required density for experiments in assay medium (RPMI (without phenol red) containing 1 % FCS, 0.5 % DMF and L-glutamine).

### Transmission electron microscopy

Cell suspensions, fixed in 2.5% glutaraldehyde, were sedimented and embedded in low melting agarose (2% in PBS). Drops of agarose containing cells were then postfixed in 0,5% osmiumtetroxide, contrasted with tannic acid and 2% uranyl acetate, dehydrated in a graded ethanol series and embedded in epoxy resin. After polymerization, sections were cut at 60 nm and contrasted with lead citrate. Specimens were analyzed in a Leo 906E transmission electron microscope at 100KV (Zeiss, Oberkochen, DE) using a side mounted digital camera (Morada; SIS-Olympus Münster DE).

### Analysis of ROS production from cells

Cells were seeded at 10^5^/well of a white 96 well plate (Nunc) in triplicates in a final volume of 100 μl in assay medium. HRP (1.2 U/ml) and luminol (50 μM) were added 10 minutes after seeding and 10 minutes later cells were stimulated with 100 nM PMA. Luminescence signal was measured using a VICTOR XLight Multimode Plate Reader (Perkin Elmer).

### Analysis of cell death

Cells were seeded at 10^5^/well of a 96 well plate in duplicates or triplicates in a final volume of 100 μl. The cell impermeable DNA dye SYTOX Green (50 nM) was added and 10 minutes later cells were stimulated with 100 nM PMA or with 5 μM A23187. SYTOX Green fluorescence signal corresponding to NET formation and cell death was measured over time using excitation/emission of 485/518 nm. Alternatively, microscopy pictures were acquired using brightfield and GFP channels (for SYTOX Green) in order to visualize the morphology of dead cells.

### Analysis of surface marker expression

PLB-985 and murine *ex vivo* differentiated bone marrow cells were stained at various time points of differentiation with indicated surface marker antibodies for 20 min at 4 °C in PBS with 2% FCS, washed and analyzed by flow cytometry using a MACS Quant Analyzer (Miltenyi Biotec). DAPI was added just before acquisition to exclude dead cells.

Murine blood was stained the same way as described above, but after staining, samples were incubated with eBioscience 1 Step Fix/Lyse Solution (invitrogen, 00-5333-54) for 1h to lyse erythrocytes before acquisition.

### CRISPR/Cas9 screen

PLB-985 were transduced with a lentiviral library containing CRISPR/Cas9 sgRNAs^21^ at an estimated MOI of <1. Cells were selected with puromycin (Sigma, 2.5 μg/ml), differentiation was performed as described for wt PLB-985. At d7 of differentiation, cells were stimulated with 100 nM PMA for 16h and surviving cells were sorted by FACS. DNA extraction was performed by DNeasy Blood & Tissue Kit (Qiagen) and PCR amplification was performed as described^21^. Reads were mapped to the library and abundance of sgRNAs was calculated per condition. Genes with greater than 50 % of their sgRNAs identified at values of at least 2-fold overrepresentation in the PMA-treated group versus the control group were defined as hits.

### Generation of CRISPR/Cas9 knockout lines

PLB-985 were transduced with lentiviral particles containing the sgRNA of interest cloned into lentiCRISPRv2, cells were selected with puromycin (2.5 μg/ml) and, if single clones were analyzed, seeded at a density of 0.8 cells/well in 96 wells for limited dilution cloning. Clones were sequenced using the OutKnocker protocol and software^24^ (Supplementary Table 3). Clones with out of frame indels leading to efficient gene disruption were used for subsequent experiments.

For GATA-1 and GATA-2 disruption in scr., H1.2 and H1.4-deficient cells, sgRNAs targeting GATA were cloned into a modified lentiCRISPRv2, in which the Cas9 sequence was exchanged for a blasticidin resistance cassette. After transduction, cells were selected with blasticidin (50 μg/ml) and used in batch populations for experiments.

### RNA extraction and qRT-PCR

Cell samples were pelleted, resuspended and lysed in TRIzol Reagent (life technologies). RNA was isolated by chloroform extraction and isopropanol precipitation. Similar amounts of total RNA were used for RT-PCR (High Capacity cDNA Reverse Transcription Kit, 4368814, Thermo Fisher Scientific). qRT-PCR was performed using SYBR Green (4385612, Thermo Fisher Scientific) on a QuantStudio 3(Thermo Fisher Scientific).

### RNA-sequencing

Total RNA of PLB-985 at various time points was isolated using RNeasy kit (Qiagen). Library preparation and illumina sequencing were performed by the Max Planck-Genome-centre Cologne, Germany (https://mpgc.mpipz.mpg.de). The data were mapped to hg38.87 using STAR 2.5.2b^25^ and differential expression analysis was performed using edgeR^26^. Gene level read counts from primary human cells at various steps of neutrophil differentiation were retrieved from Blueprint^27^ (http://www.blueprint-epigenome.eu, Datasets EGAD00001002446, EGAX00001244028, EGAX00001244022 and EGAD00001002366) and analyzed the same way. Gene module enrichment was performed using tmod package^28^ and concordance and discordance was analyzed by calculating discordance scores^29^. Eosinophil gene sets were retrieved from Harmonizome^30^ and Broad Institute (http://software.broadinstitute.org/gsea, Nakajima set), GATA-1 gene set was generated by associating GATA-1 ChIP seq peaks (GEO accession GSM1003608) to genes using ChIPseeker^31^.

### Ex vivo differentiation of murine bone marrow cells into neutrophils and eosinophils

Murine bone marrow was extracted from the femurs and tibiae of adult age-matched mice. Cells were flushed, erythrocytes were lysed by osmotic lysis, cells were washed and lineage-negative precursor cells were sorted by negative selection using magnetic beads (130-110-470, Miltenyi Biotec). Lineage-negative cells were subsequently propagated analogous to the protocol described^32^. Briefly, cells were seeded in IMDM supplemented with 20% FCS, antibiotics, L-glutamine and 0.1 μM 2-mercaptoethanol (Sigma), as well as with the recombinant cytokines IL-3 (20 ng/ml), IL-5 (50ng/ml), IL-9 (50 ng/ml), GM-CSF (10 ng/ml) and SCF (20 ng/ml) for 6 days.

### Statistical analysis

The descriptive statistics of all figure panels are mentioned in the respective figure legends. Populations of circulating leukocytes in murine blood were compared by a two-tailed, unpaired t test and p values are indicated above the populations. RNA-seq data were analyzed as described in the respective methods section.

## Results

### PLB-985: a model for neutrophil maturation and function

We used the human promyelocytic cell line PLB-985 ^33^. PLB-985 differentiate into neutrophil-like cells that resemble primary human neutrophils in various ways^20, 34–36^. PLB-985 changed their nuclear morphology and developed granules, though fewer than primary neutrophils, throughout the 7 days of differentiation (referred as d0 to d7 in this report) (Fig. 1a). Once differentiated, PLB-985 also produced ROS in response to the agonist phorbol 12-myristate 13-acetate (PMA) (Fig. 1b, Supplementary Fig. 1a) and upregulated the surface marker CD11b (Fig. 1c). Furthermore, differentiated PLB-985 phagocytosed *Escherichia.coli*, a process that could be blocked by the phagocytosis inhibitor cytochalasin B (Supplementary Fig. 1b). Importantly, PMA potently induced cell death in fully differentiated PLB-985, but not in d0 or d3 cells (Fig. 1d). PMA-induced cell death was morphologically similar to NETosis; dying PLB-985 expanded their nuclei and a significant fraction released DNA as shown by SYTOX Green staining (Fig. 1e, Supplementary Fig. 1c). Of note, the calcium ionophore A23187, another stimulant of NET formation, also induced cell death in PLB-985, both at d3 and d7 of differentiation (Supplementary Fig. 1d). Taken together, these experiments show that PLB-985 cells differentiate into neutrophil-like phagocytes that produce ROS and NETs.

**Fig. 1.**
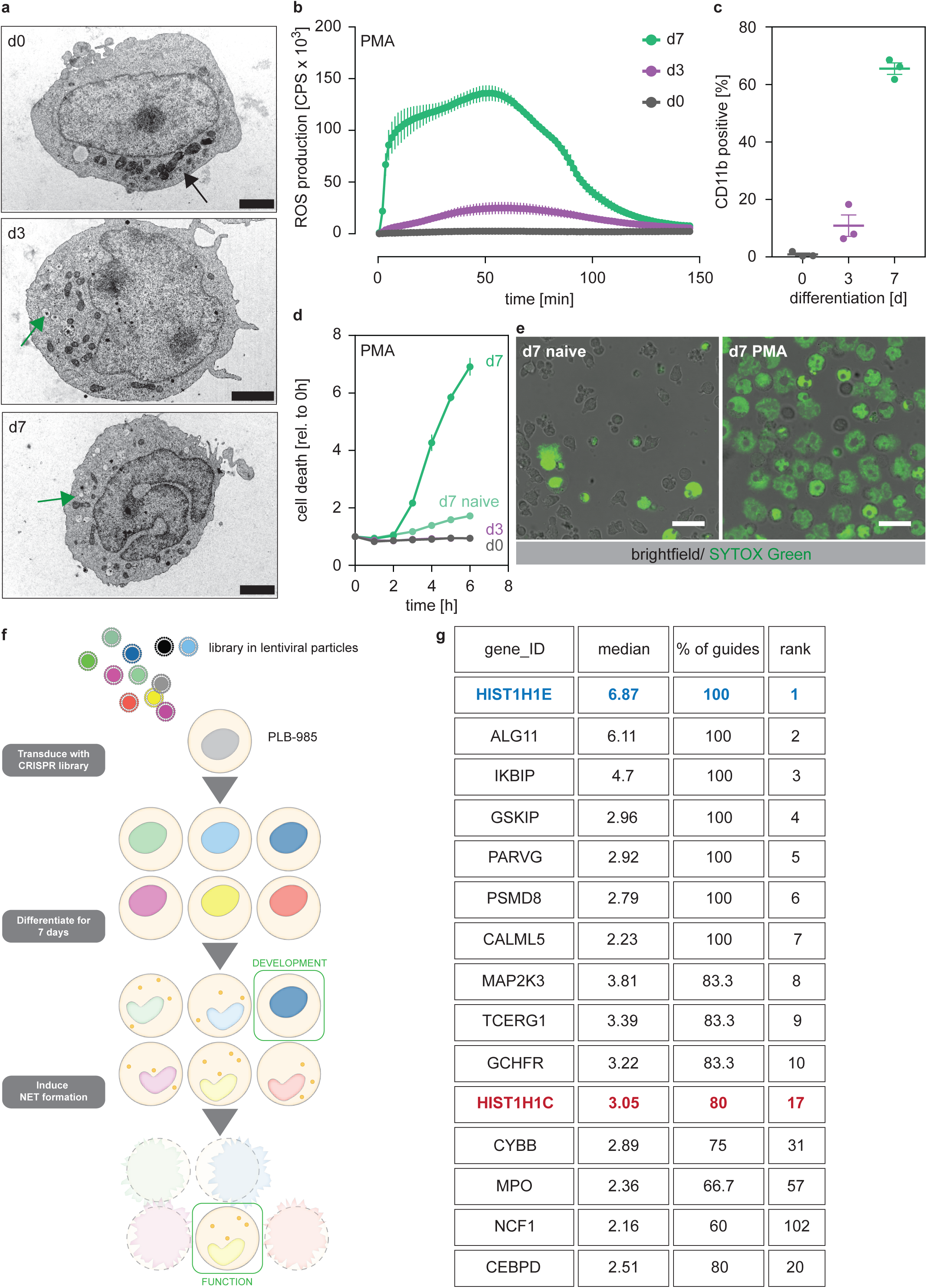
A genome-wide screen to identify genes required for PLB-985 cells differentiation and function. **a-d** Characterization of PLB-985 differentiation and function at d0, d3 and 7 of differentiation. **a** Electron microscopy images of PLB-985 showing acquisition of granules (green arrows, the black arrow at d0 indicates mitochondria) and changes in nuclear morphology. Scale bars correspond to 5 μm. **b** ROS production of PLB-985 in response to 100 nM PMA, showing that only fully differentiated PLB-985 produced an oxidative burst. **c** Surface expression of CD11b, depicted is the percentage of CD11b positive cells out of all viable singlets. **d** PMA-induced cell death was measured over time after addition of the cell-impermeable DNA dye SYTOX Green and analysis of fluorescence indicating cell death. “d7 naïve” indicates differentiated cells, which were not treated with PMA. **e** Representative images of differentiated PLB-985 (d7) after addition of SYTOX Green and treatment with or without PMA. Scale bars are 20 μm. **f** Outline of the CRISPR/Cas9 screen. Cells were transduced with a genome-wide CRISPR/Cas9 library (lentiviral particles, small circles), differentiated for 7 days, treated with PMA for 16h and survivors were sorted and sequenced to identify sgRNAs. **g** Top 10 of the screen, H1.2 and indicated genes with known neutrophil functions, ranked first by % of overrepresented guides and then by median overrepresentation and mapped to a proteome list of human primary neutrophils. **b-d** Depicted are mean -/+ SEM of 3 independent experiments.

To show that PLB-985 can be genetically modified by CRISPR/Cas9, we disrupted expression of NOX2 (*CYBB*), a component of the NADPH oxidase complex, which is required for PMA-induced NETosis^37^. We also transduced PLB-985 with a non-target scrambled single-guide RNAs (sgRNA) as a control (subsequently called scr). Differentiated PLB-985 cells lacking NOX2 failed to produce an oxidative burst and did not undergo cell death in response to PMA (Supplementary Fig. 1e, f). To inactivate NADPH oxidase independently of NOX2, we also disrupted expression of its NCF2 subunit, and showed again that these cells did not die upon PMA stimulation (Supplementary Fig.1g). These results demonstrate that PLB-985, which can be genetically modified, are a suitable model to mimic human neutrophils.

### A genome-wide screen to identify regulators of neutrophil maturation and function

Since we found that PMA-induced NET formation only occurred in fully differentiated PLB-985, we used this readout to perform a genome-wide CRISPR/Cas9 screen to identify genes required for neutrophil maturation and function (Fig. 1f). We reasoned that, besides NET-defective cells, we could also identify differentiation-defective cells by sorting cells that survived PMA treatment. We transduced PLB-985 with a library targeting all human genes with 5-6 sgRNAs per gene^21^. As expected, PMA stimulation reduced the fraction of identified guides in the survivor population (Supplementary Fig. 1h), indicating selection of specific clones.

We defined “hits” as genes for which at least 50 % of the identified sgRNAs were overrepresented (more than two-fold) in our survivor population. Underlining the validity of the screening approach, we identified NOX2, NCF1 and MPO as hits; all three are known to be required for PMA-induced NET formation^37, 38^ (Fig. 1g, Supplementary Table 1). We also found C/EBP-δ, a member of the C/EBP family of transcription factors that are crucial for granulopoiesis^39^ (Fig. 1g). Unexpectedly, we identified two members of the linker histone H1 family among the 20 highest ranked genes. *HIST1H1E* (H1.4) was overrepresented with 100 % of sgRNAs and showed the highest median enrichment (Fig. 1g, Supplementary Table 1). Notably, there are no studies showing a functional role of H1 in neutrophil differentiation or NET formation.

### H1.4 and H1.2 affect neutrophil function through differentiation

There are five somatic replication-dependent (H1.1 – H1.5) and two replication-independent H1 subtypes (H1.0 and H1X). PLB-985 expressed H1.2 and H1.4 more abundantly than the other subtypes (Fig. 2a). The expression of replication-dependent H1 subtypes occurs mainly in the S phase of proliferating cells and, accordingly, H1 mRNA levels decreased during PLB-985 differentiation (Supplementary Fig. 2a, same experiments as in Fig. 2a, but plotted as relative to d0). As expected, this reduction was more evident for subtypes with a higher baseline expression (Fig. 2a, Supplementary Fig. 2a).

**Fig. 2.**
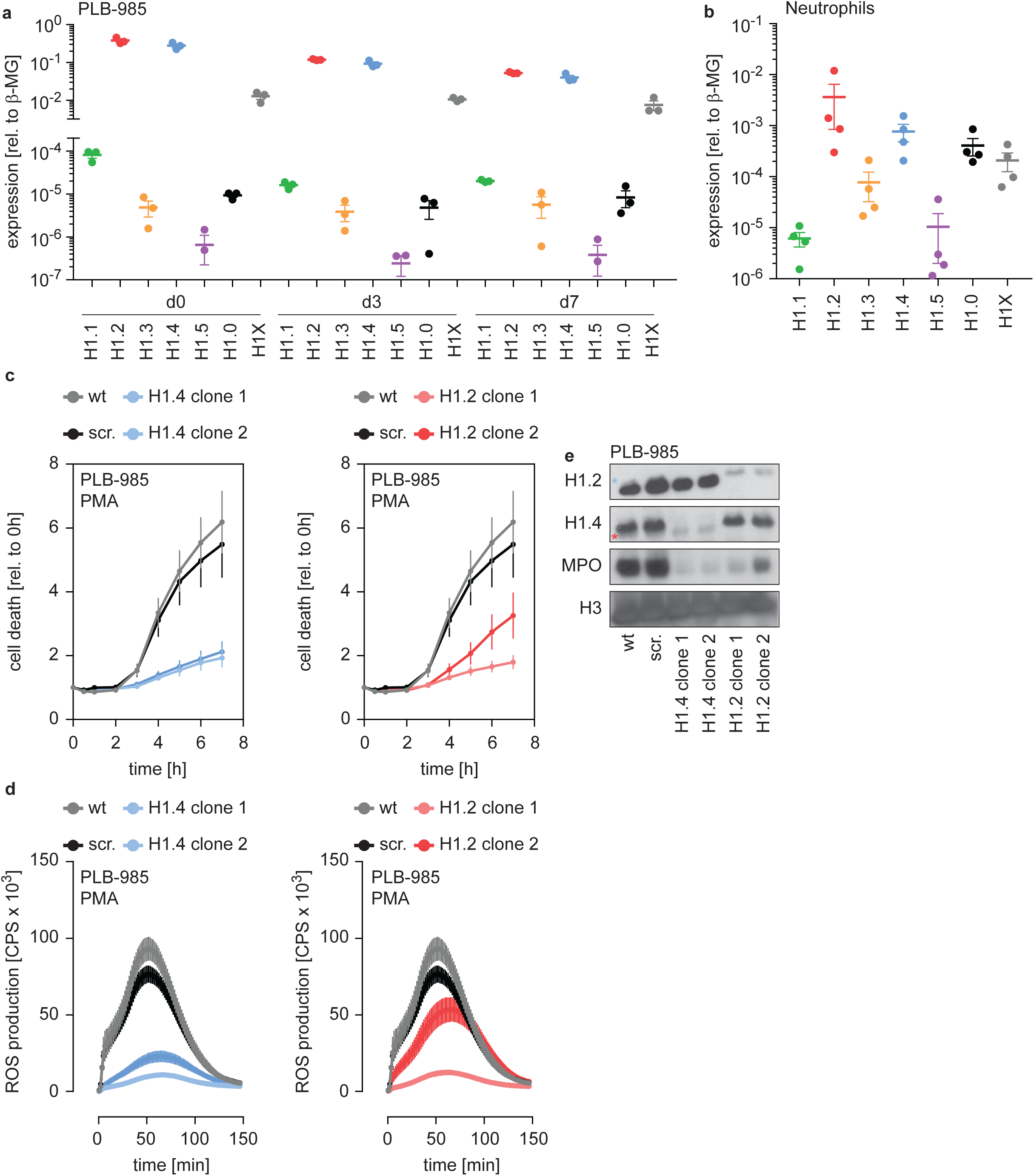
H1.2 and H1.4 are required for PLB-985 differentiation. **a** mRNA expression levels of indicated H1 subtypes in PLB-985 cells at d0, d3 and d7 of differentiation, relative to the housekeeping gene β-microglobulin, depicted is the mean -/+ SEM of 3 independent experiments. **b** mRNA expression levels of indicated H1 subtypes in human primary neutrophils, relative to β-microglobulin. Data points are from 4 different donors, error bars are mean -/ + SEM. **c** PMA-induced cell death of wild type (wt) scrambled sgRNA (non-target, scr.) and two clones of either H1.2 or H1.4 over time, measured by SYTOX Green fluorescence. Depicted are mean -/+ SEM of 5 independent experiments of cells at d7 of differentiation. **d** Measurement of PMA-induced ROS production from wt, scr. and H1.2 or H1.4 deficient PLB-985, depicted are mean -/+ SEM of 5 independent experiments at d7 of differentiation. **c, d** wt and scr. values are the same in the left and right panels, respectively. **e** Western blot of lysates of PLB-985 at d3 of differentiation, showing efficient disruption of H1.2 and H1.4 (both antibodies recognize the other subtype, which is marked by a red asterisk for H1.2 and a blue asterisk for H1.4) as well as reduced MPO expression, the core histone H3 served as loading control.

We also analyzed a dataset comparing mRNA expression of human primary bone marrow cells undergoing differentiation into neutrophils at various stages of maturation, namely myelocytes, metamyelocytes, band form neutrophils and segmented neutrophils^27^ (see methods section for RNA-seq). Consistent with our PLB-985 data, we found a similar reduction of H1 mRNA as maturation progressed (Supplementary Fig. 2b).

mRNA levels of each H1 subtypes were still detectable in differentiated PLB-985 and primary cells. Accordingly, human primary neutrophils isolated from blood of healthy donors expressed detectable amounts of the H1 genes. H1.2 and H1.4, the two hits in our genome wide screen, were the most abundantly expressed subtypes (Fig. 2b). Interestingly, peripheral blood mononuclear cells (PBMCs) and monocytes showed a different expression pattern of this gene family than neutrophils (Supplementary Fig. 2c, d).

To verify the results from our screen, we disrupted expression of H1.2 and H1.4 in PLB-985 by targeting these genes with CRISPR/Cas9. We used two clones deficient for H1.2 (from 2 sgRNAs) and two clones deficient for H1.4 (from 1 sgRNA) in our analysis. Cell death was reduced in both H1.2 and H1.4-deficient clones, confirming the results of the screen (Fig. 2c). H1.2 and H1.4-deficient clones produced ROS less efficiently than wild type cells upon PMA stimulation (Fig. 2d). Furthermore, cells deficient for either of the two H1 subtypes expressed less MPO (Fig. 2e). These findings suggest that H1.2 and H1.4 are required for PLB-985 to differentiate into mature, neutrophil-like cells.

### Opposing effects of H1 subtypes on neutrophil differentiation

To confirm that deficiency of H1.2 and H1.4 affected PLB-985 maturation, we showed that surface expression of the differentiation marker CD11b is decreased on both d3 and d7 of differentiation (Fig. 3a, b, Supplementary Fig. 3a, b) in cells where these genes were deleted. We also disrupted expression of the other three somatic H1 subtypes. Surprisingly, loss of these subtypes resulted in the opposite phenotype. We observed markedly enhanced expression of CD11b already at d3, especially in clones deficient in H1.1 and H1.5 (Fig. 3a, b, Supplementary Fig. 3a, b).

**Fig. 3.**
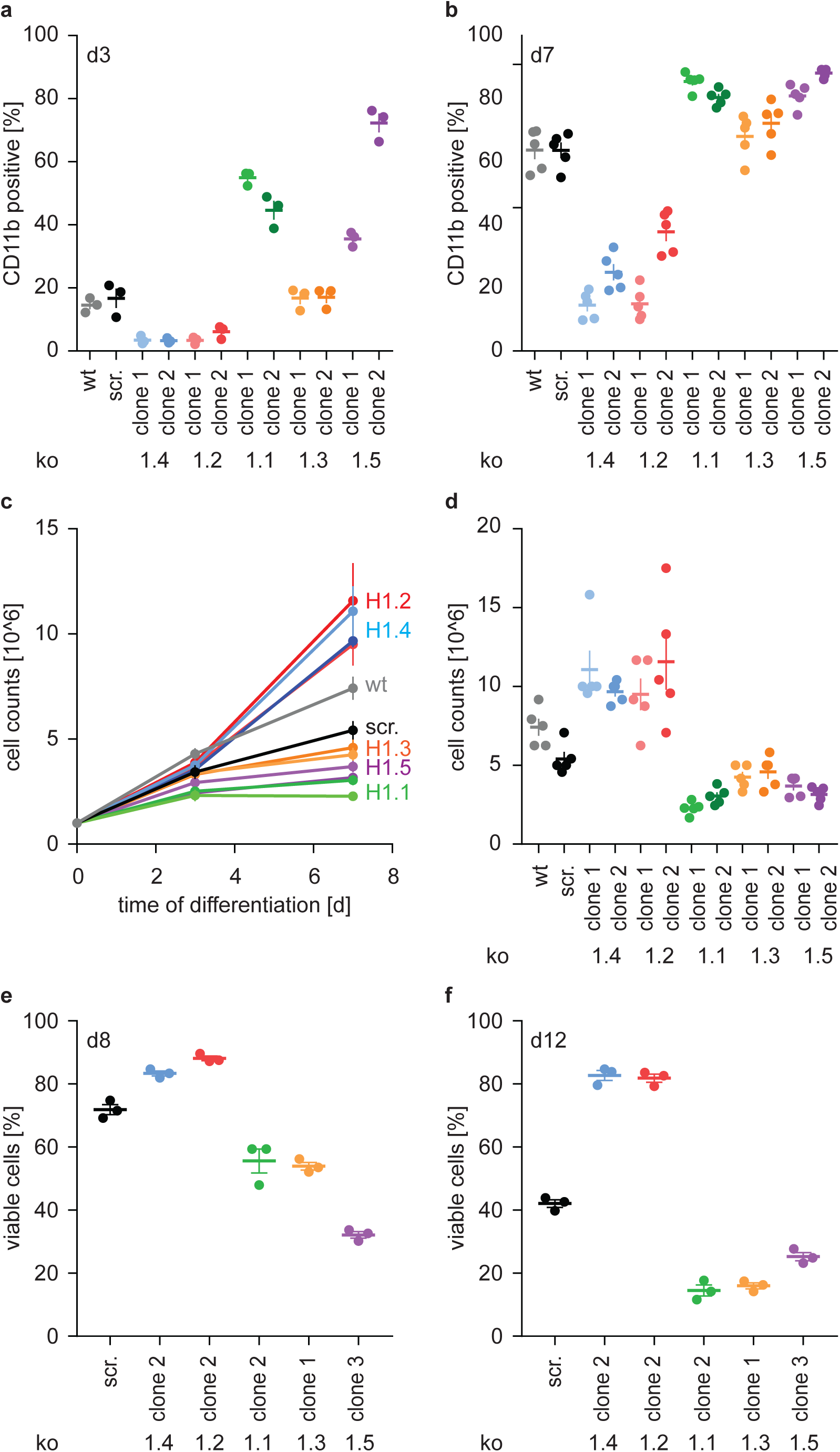
H1 effects on PLB-985 differentiation are subtype-specific. **a, b** Expression of the surface marker CD11b at d3 (**a**) or d7 (**b**) of differentiation. Depicted is the percentage of CD11b-positive cells out of all living singlets for wt, scrambled (scr.) and two knock-out clones per H1 subtype. **c** Growth curves of PLB-985 during differentiation and (**d**) cell counts at d7 of differentiation show restricted growth of H1.1, H1.3 and H1.5-deficient cells and enhanced growth of H1.2 and H1.4-deficient clones. **a-d** Depicted are 5 independent experiments, error bars are SEM. **e, f** Viability, as determined by DAPI negative cells, at d8 (**e**) and d12 (**f**) of differentiation of 3 independent experiments. Error bars are SEM.

Differentiation of neutrophils and PLB-985 is terminal - if not activated, these cells die by apoptosis. The number of wild type and control (scr.) PLB-985 cells increased more from d0 to d3 than from d3 to d7. This suggests that cells withdrew from proliferation (Fig. 3c). Interestingly, H1.2 and H1.4-deficient clones grew unrestrictedly until d7, whereas cells deficient in H1.1, H1.3 and H1.5 decreased in comparison to controls (Fig. 3c, d). Furthermore, control cells successively lost viability from d8 to d12 (Fig. 3e, f). In line with their enhanced growth, H1.2 and H1.4-deficient cells were more viable than controls, whereas H1.1, H1.3 and H1.5-deficient cells showed less viability than controls (Fig. 3e, f). Of note, cells deficient in H1.1 had a stronger oxidative burst in response to PMA than control cells, consistent with enhanced maturation (Supplementary Fig. 3c). Clones deficient in H1.3 and H1.5, on the other hand, produced less ROS than control cells (Supplementary Fig. 3d, e). This could be a specific effect of H1.3 and H1.5, but it could also reflect reduced viability at the onset of the experiment. In summary, these experiments show an unexpected and opposing effect of H1 subtypes on PLB-985 cell differentiation and function.

### The impact of H1 deficiency manifests before onset of differentiation

To understand the extent of H1 subtype-specific effects on neutrophil differentiation we performed RNA-sequencing of the knockout lines. We sampled wild type cells, scr. cells and two clones per H1 subtype at d0, d3 and d7 of differentiation in 4 independent experiments (Supplementary Table 2). To validate the applicability of our experiment, we compared the transcriptome of control PLB-985 cells (the combination of wild type and scr. samples) to human primary cells at various differentiation stages towards neutrophils (also see methods section). Gene set enrichment analysis demonstrated that PLB-985 and primary cells concordantly regulate processes during differentiation (Fig. 4a, Supplementary Fig. 4a, b. Supplementary Fig. 4a depicts the same analysis as Fig. 4a with the gene set labels). Using principal component analysis (PCA), we saw that these cells follow a similar differentiation trajectory (Supplementary Fig. 4c). As expected, among the most robustly upregulated genes during differentiation of control cells we found several with known functions in neutrophils (Fig. 4b).

**Fig. 4.**
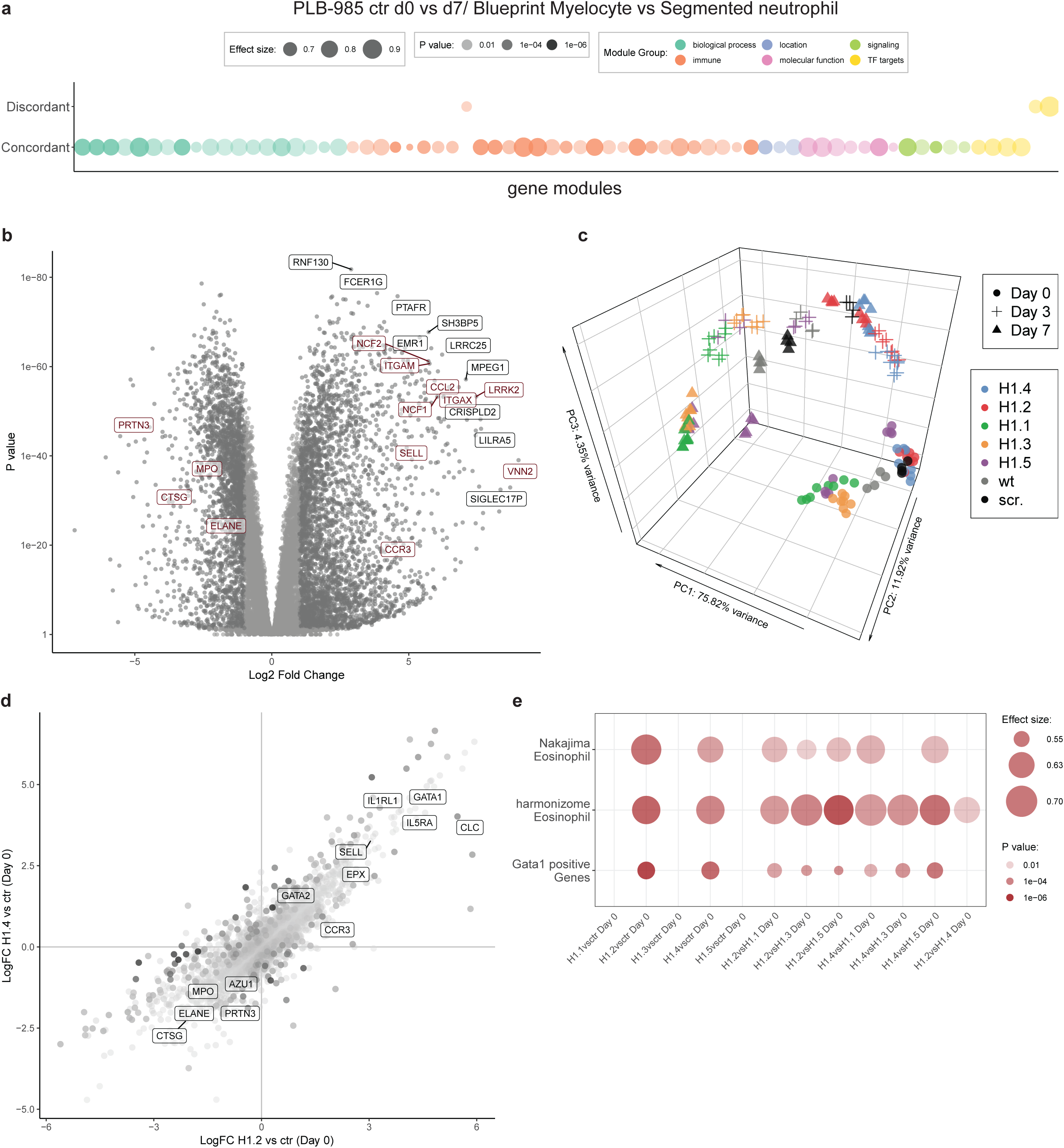
RNA-seq reveals H1 subtype-specific effects on differentiation and an eosinophilic signature of undifferentiated H1.2/H1.4 knockout lines. **a** Concordantly and discordantly regulated genes when comparing PLB-985 differentiation (wt and scr., d0 to d7) and human neutrophil differentiation (Blueprint, neutrophilic myelocyte to segmented neutrophil of bone marrow) using discordance scores and tmod. Most modules are concordantly regulated. **b** Volcano plot of all differentially expressed genes of control cells (wt and scr.) between d7and d0, darker gray depicts p<0.05 and logFC >2. A selection of strongly upregulated genes is labelled and neutrophil markers are highlighted in red. Expression of granule proteins, such as MPO, neutrophil elastase (ELANE), proteinase 3 (PRTN3) and cathepsin G (CTSG) decreases during differentiation. **c** Principal component analysis (PCA) of control and mutant PLB-985 during differentiation shows clustering according to experimental replicates and genotypes and separates samples according to differentiation state. While lines deficient in H1.1, H1.3 and H1.5 already at d3 cluster closely to wt and scr. at d7, indicating faster and more efficient differentiation, H1.2 and H1.4-deficient clones at d7 still only cluster with wt and scr. at d3, indicating a stalled development. **d** Scatter dot plot depicting differentially expressed genes of H1.2 knockouts vs control cells at d0 of differentiation (x axis) and H1.4 knockouts vs control cells at d0 of differentiation (y axis), color (alpha) indicates p-value of differential expression between H1.2 and H1.4 at d0. Neutrophil genes such as neutrophil elastase (ELANE), proteinase 3 (PRTN3), cathepsin G (CTSG), azurocidin (AZU1) and MPO are strongly downregulated in both conditions when compared to wt and scr., whereas eosinophil genes such as galectin 10 (CLC), interleukin-5 receptor alpha (IL5Ra), GATA-1, GATA-2 or eosinophil peroxidase (EPX) are upregulated. **e** Gene set enrichment analysis of indicated samples at d0 of differentiation shows enhanced expression of eosinophil and GATA-1 gene sets (see methods) in H1.2 and H1.4-deficient lines, but not in other knockout conditions.

We analyzed the H1-deficient clones by PCA and found that H1.2 and H1.4-deficient clones at d7 clustered more closely with control cells at d3, indicating delayed differentiation (Fig. 4c). H1.1, H1.3 and H1.5-deficient cells at d3 clustered with control cells at d7, suggesting accelerated differentiation (Fig. 4c).

Analysis of the enrichment of gene modules confirmed that, during differentiation, control cells upregulated genes related to the innate immune response and downregulated genes corresponding to cell cycle and division, in line with the context of terminal differentiation (Supplementary Fig. 4d). Interestingly, in H1.2 and H1.4-deficient cells, the modules were less regulated, again demonstrating prolonged proliferation and obstruction of differentiation (Supplementary Fig. 4d). We performed the same analysis for clones deficient in H1.1, H1.3 and H1.5 and confirmed accelerated differentiation (Supplementary Fig. 4d). We subsequently asked whether absence of H1 subtypes also affected the determination and lineage choice of PLB-985 before differentiation. To address this question, we compared differentially expressed genes of H1.2 and H1.4-deficient cells and control cells at d0. We found that both H1.2 and H1.4-deficient cells downregulated neutrophil genes, such as MPO and neutrophil proteases among others (Fig. 4d), indicating that lineage fate was affected in undifferentiated cells.

Surprisingly, among the upregulated genes we found a significant number of genes associated with eosinophils and a GATA-1 signature (Fig. 4d, e, see methods section for the source of datasets) We found this upregulation specifically in clones deficient in H1.2 and H1.4 in comparison to all other genotypes (Fig. 4e). This suggests that loss of H1.2 and H1.4 affects lineage determination of PLB-985 cells even before the onset of differentiation by driving them towards a more eosinophil-like fate. These data indicate that H1 subtype depletion affects both cell fate determination and differentiation per se.

### Enhanced eosinophil differentiation of H1.2/H1.4-deficient hematopoietic stem cells

Loss of H1.2 and H1.4 appears to bias the lineage commitment of PLB-985 away from neutrophils via regulation of an eosinophil-like transcriptional program. To test whether this bias also exists in primary cells, we analyzed cells isolated from H1.2/H1.4 double-deficient mice^13^. The profile of circulating leukocytes in the blood of adult animals was indistinguishable from wild type animals (Supplementary Fig. 5a-e). Furthermore, wild type and H1.2/H1.4-deficient animals had the same proportion of neutrophils in circulation (Fig. 5a). We isolated lineage negative cells from murine bone marrow and differentiated them into several lineages^31^. Importantly, H1.2/H1.4-deficient cells expressed elevated levels of surface markers for eosinophils and decreased levels of neutrophil surface markers, as compared to wild type cells. This recapitulates the phenotype we observed in PLB-985 (Fig. 5b-d). These experiments show that H1.2 and H1.4 cooperate to determine the neutrophil-eosinophil cell fate decision in hematopoietic progenitor cells. Furthermore, this suggests that, *in vivo,* the H1.2 and H1.4 deficiency is compensated to generate normal amounts of neutrophils during homeostasis.

**Fig. 5.**
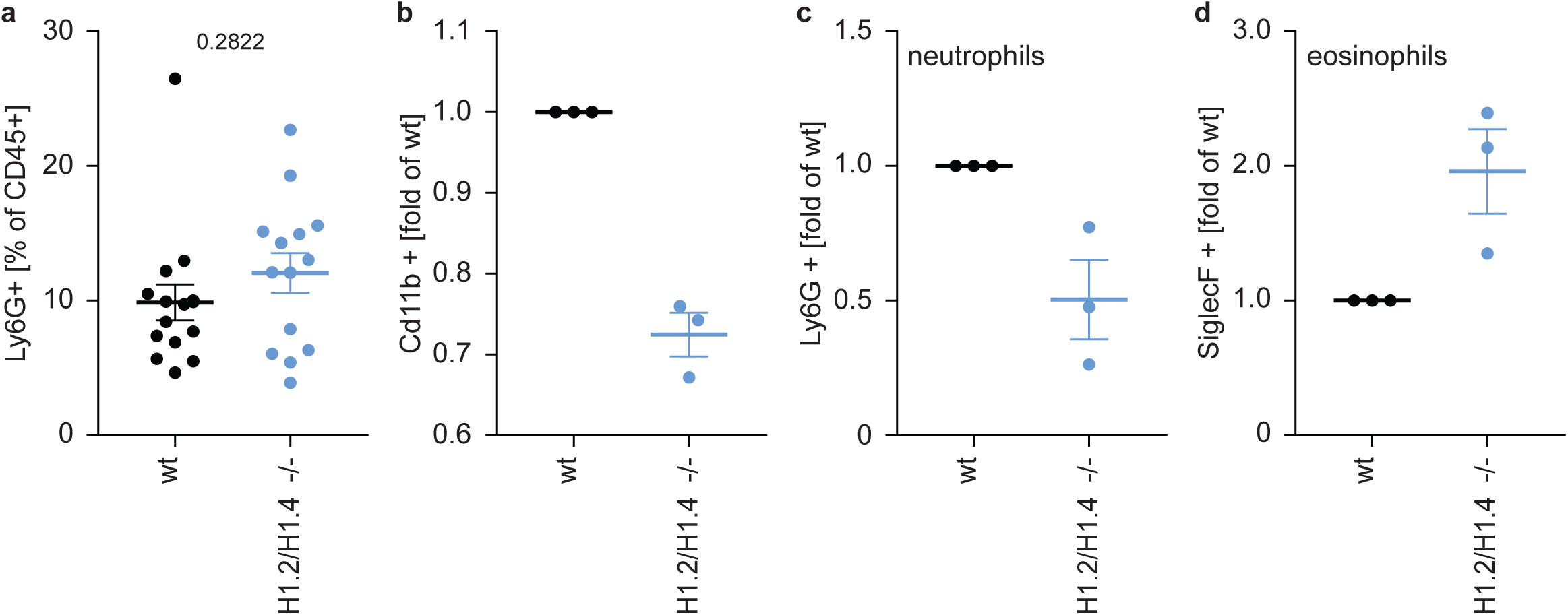
Deletion of H1.2 or H1.4 biases murine hematopoietic stem cells towards an eosinophil lineage-. **a** Analysis of circulating neutrophils in wild type (wt, n=15) and H1.2/H1.4-double deficient mice (H1.2/H1.4 -/-, n=14). Flow cytometry analysis of neutrophils in whole blood of adult age-matched animals, other immune cell types are shown in Supplementary Fig. 5. **b-d** Lineage-negative hematopoietic stem cells were sorted from murine bone marrow and cultured for 6 days in the presence of various cytokines allowing differentiation into several immune cell lineages. Depicted are CD11b positive cells (**b**) and the fraction of neutrophils (**c**) or eosinophils (**d**) within CD11b-positive cells. Each dot represents an independent experiment consisting of two mice per genotype, the mean of the two wild type animals was set to 1 and the mean of the two H1.2/H1.4 -/- animals is depicted relative to wt.

### Loss of H1.2 and H1.4 affect neutrophil cell fate determination via GATA-2

Two transcription factors required for eosinophil differentiation, GATA-1 and GATA-2, were upregulated in H1.2 and H1.4 knockout lines (Fig. 4e). GATA-1 transcripts were very low in all lines sequenced (Supplementary Fig. 6a). GATA-2 transcript levels at d0 inversely correlated with the potential of PLB-985 to adopt a neutrophil-like cell fate (Supplementary Fig. 6b). We speculated that H1 subtypes affect neutrophil differentiation through either GATA-1 or GATA-2. To test this hypothesis, we disrupted expression of GATA-1 and GATA-2 by CRISPR/Cas9 and analyzed batch populations of various mutations at d4 of differentiation. We chose d4 because we assumed that cells still expressing GATA could outgrow GATA-deficient cells at later time points. Interestingly, GATA-2 deficiency enhanced maturation of PLB-985, as indicated by an increased ability to produce ROS in response to PMA (Supplementary Fig. 6c). In contrast, GATA-1 knockout lines behaved as scr. cells (Supplementary Fig. 6c).

**Fig. 6.**
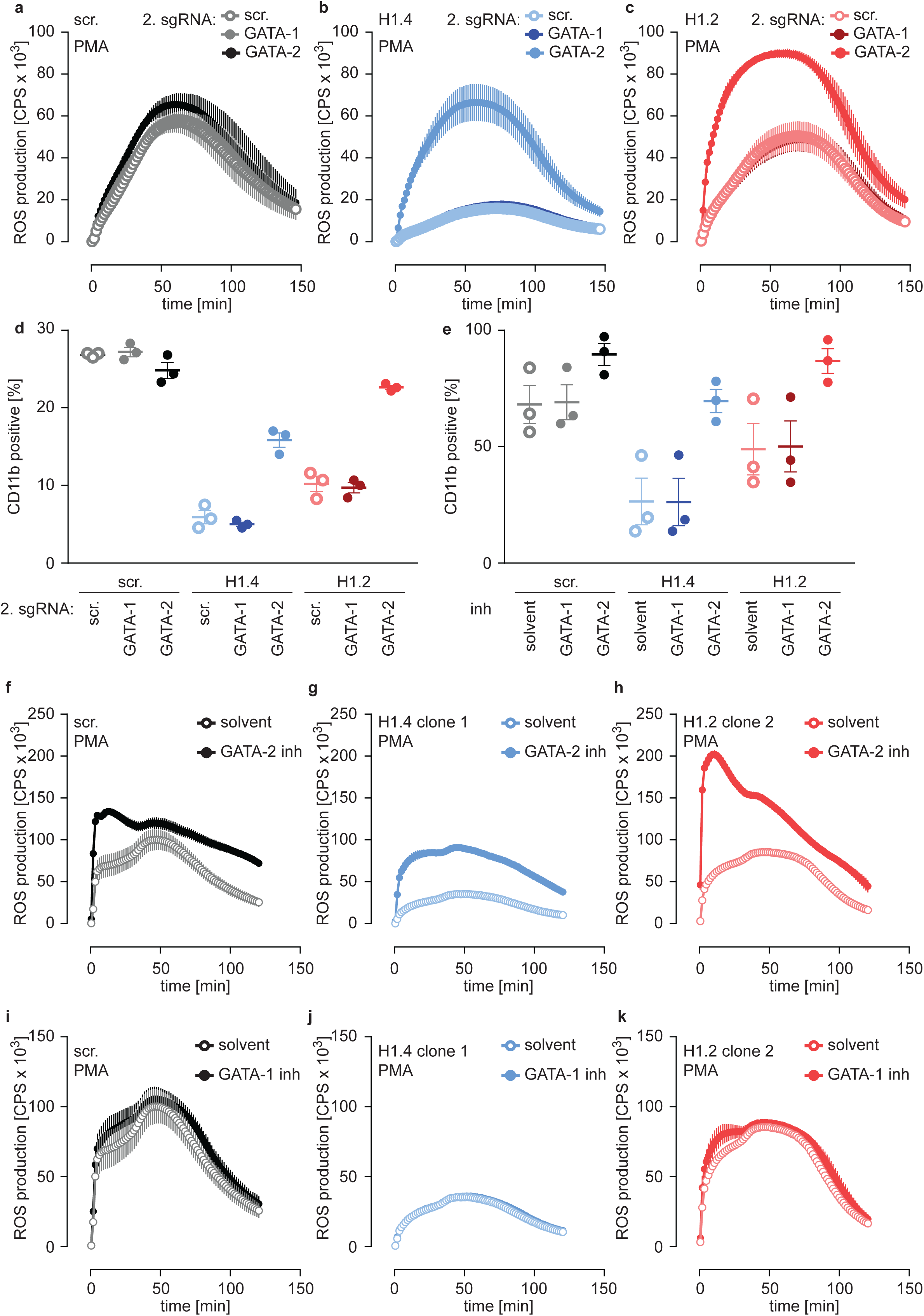
Deficiency of H1.2 or H1.4 is rescued by inhibition of GATA-2. **a-c** Deficiency in GATA-2 but not GATA-1 rescues H1.2 and H1.4 deficiency. Populations are batches of PLB-985. Scr. (**a**), H1.4 (**b**) and H1.2 (**c**) knockout lines were transduced with an sgRNA against GATA-1 or GATA-2 or with a scr. sgRNA as control and analyzed for ROS production in response to PMA at d4 of differentiation. **d** Deficiency in GATA-2 rescues CD11b expression of H1.2 and H1.4-deficient PLB-985. CD11b expression of the indicated genotypes was measured at d4 of differentiation. **e** Treatment of PLB-985 with GATA-2 but not GATA-1 inhibitor rescues expression of CD11b, especially in lines deficient in H1.2 and H1.4. **f-k** ROS production of PLB-985 in response to PMA is affected upon GATA-2, but not GATA-1 inhibition during differentiation. **f-k** PLB-985 ROS production was measured at d7 of differentiation, indicated genotypes were treated with indicated inhibitors or solvent before the onset of differentiation and at d4. **a-k** Shown is the mean -/+ SEM of 3 independent experiments.

To test whether the differentiation phenotype of H1.2 and H1.4-deficient lines relied on GATA-2, we performed a rescue experiment by generating double-deficient lines. We transduced scr. cells and H1.2 or H1.4-deficient cells with GATA-1 and GATA-2 sgRNAs or with a scr. sgRNA as a control and analyzed these populations at d4. Importantly, disruption of GATA-2 expression rescued ROS production (Fig. 6a-c) and CD11b expression (Fig. 6d) in H1.2 and H1.4-deficient PLB-985. Again, disruption of GATA-1 had no effect (Fig. 6a-d). These experiments demonstrate that the differentiation phenotype resulting from H1.2 and H1.4 deficiency depends on upregulation of GATA-2 and, importantly, that loss of GATA-2 is sufficient to allow differentiation in the absence of these H1 subtypes.

In a second approach, we treated PLB-985 with inhibitors for GATA-1 and GATA-2, starting treatment one day before differentiation. Inhibition of GATA-2 enhanced CD11b expression of H1.2 or H1.4-deficient lines (Fig. 6e). To confirm the effect of GATA-2 inhibition on differentiation, we measured mRNA expression of two genes, one that increased (aquaporin-9, AQP9) and one that decreased (MPO) during differentiation (Supplementary Table 2). As expected, H1.2 and H1.4-deficient lines expressed less AQP9 and more MPO than scr. cells at d7 of differentiation (Supplementary Fig. 6d, e). GATA-2 inhibition upregulated expression of AQP9 and downregulated expression of MPO in H1.2 and H1.4 knockout lines at d7 of differentiation, indicating a rescue of maturation (Supplementary Fig. 6d, e). In line with our results with CRISPR/Cas9-mediated gene disruption, GATA-2 inhibition restored the capacity of H1.2 and H1.4-deficient clones to mount an oxidative burst (Fig. 6f-h). As expected, treatment with the GATA-1 inhibitor did not affect maturation and function of PLB-985 of either genotype (Fig. 6e, i-k). The combination of GATA-1 and GATA-2 inhibitors did not have an additive effect on PLB-985 maturation, confirming that GATA-2 is the relevant transcription factor for the phenotype observed in H1.2 and H1.4-deficient clones (Supplementary Fig. 6f). Importantly, the response to either GATA-2 deficiency or inhibition was stronger in H1-deficient PLB-985 than in scr. cells. This is in line with enhanced baseline expression of the transcription factor in H1.2 or H1.4-deficient clones. These data show that H1.2 and H1.4 affect fate determination and subsequent neutrophil differentiation and function at least partly through the regulation of GATA-2.

## Discussion

We identified an unexpected role of H1 subtypes in neutrophil development; loss of H1.2 and H1.4 inhibits differentiation while deficiency in H1.1, H1.3 and H1.5 enhances it. We propose that in neutrophil precursors, H1 subtypes control the expression or activity of GATA-2, which determines cell fate via transcription of neutrophil or eosinophil-specific genes. Enhanced expression of eosinophil genes – or reduced expression of neutrophil-specific genes – disrupts normal maturation, leading to prolonged proliferation and viability and reduced neutrophil function. Consistent with this, genetic ablation or inhibition of GATA-2, but not GATA-1, rescued the differentiation of H1.2 and H1.4-deficient cells. Interestingly, another study found that overexpression of GATA-2 in murine progenitor cells switched their lineage determination from macrophages to a megakaryocyte state^40^. GATA-2 therefore acts as a key molecule for fate determination of immune cells.

Importantly, we observed a preference for eosinophil development in hematopoietic stem cells isolated from the bone marrow of H1.2/H1.4-deficient mice. However, adult H1.2/H1.4-deficient animals had normal numbers of circulating neutrophils. It is possible that the turnover rate or the longevity of neutrophils is affected in these mice. Indeed, neutropenia in peripheral tissues triggers *de novo* production of neutrophils in the bone marrow^41^, demonstrating that granulopoiesis *in vivo* is regulated by feedback loops. This suggests that initial neutrophil deficiency in adult H1.2/H1.4-deficient mice could be compensated by a different cytokine milieu. Interestingly, in human breast cancer cells, combined loss of H1.2 and H1.4 leads to an upregulation of an interferon signature^42^. Although we did not observe an obvious interferon signature in H1.2 or H1.4-deficient PLB-985, it is possible that in H1.2/H1.4-deficient mice some cell types contribute to immune cell differentiation via interferon-stimulated genes.

H1 can act in redundant and subtype-specific ways^12^. Mice lacking one or two H1 subtypes are viable and fertile^13^. Analogously, PLB-985 deficient in any of the five H1 subtypes that we disrupted were viable and did not show growth defects. This is in contrast to another study, where shRNA-mediated knock down of H1.2 and H1.4 reduced cell growth and viability in various non-immune cell lines^16^. We found that loss of H1.2 and H1.4, in the context of neutrophil differentiation, enhanced viability, demonstrating cell type specific effects of H1 subtypes.

H1 has long been considered a general repressor of transcription^43^. Accordingly, a study in human cells found that genomic regions associated with active transcription were devoid of H1 subtypes^44^. However, murine embryonic stem cells that are triple-deficient in H1.2, H1.3 and H1.4 show a surprisingly low number of only 38 differentially expressed genes when compared to wild type cells^45^. Our data show that loss of H1 subtypes in more differentiated cells affects the expression of many genes with clear functional consequences. We postulate that only a fraction of differentially expressed genes in PLB-985 is directly regulated by H1 subtypes. The enhanced expression of lineage-determining transcription factors, such as GATA-2, leads to further changes in the transcriptome and affects cell differentiation.

An important open question is: how do H1 subtypes control expression of GATA-2? It is possible that H1 subtypes directly occupy the promoter or enhancer elements of GATA-2 and silence its expression. H1 subtype-specific binding to genomic regions has, for example, been shown for human H1.5^46^. Interestingly, H1.5 targeted over 20-fold more regions in differentiated human lung fibroblasts than in embryonic stem cells and H1.5 binding led to chromatin compaction and gene silencing. It is therefore tempting to speculate that similar mechanisms are at play during neutrophil development. In addition to a direct repressive effect of H1.5, its binding to chromatin was also associated with an enrichment of demethylation of histone H3 at lysine 9 (H3K9me2)^46^. H3K9me2 is a posttranslational modification of a core histone that is associated with a repressive chromatin state. H1 subtypes are therefore likely to regulate transcription via epigenetic marks on core histones. Furthermore, H1 subtypes themselves are posttranslationally modified in a subtype-specific manner^12^. Although less is known about H1 modifications than about core histones, it is likely that such modifications can determine subtype-specific protein-protein interactions, chromatin binding properties or other features that subsequently cause transcriptional changes.

It is puzzling that vertebrates express so many different H1 subtypes. Although they are able to compensate each other in “extreme” situations, such as the loss of one or even two subtypes in a whole organism^13^, they may also have more subtle and specific functions. This study contributes to our understanding of how different H1 subtypes initiate distinct transcriptional programs. Importantly, these programs appear to be evolutionarily conserved, since we found similar responses in human and murine cells.

In summary, we identified novel and opposing functions for specific H1 subtypes in granulopoiesis. H1 subtypes affect the lineage determination between neutrophils and eosinophils. We speculate that inappropriate expression of H1 subtypes in the hematopoietic compartment shapes pathologies with neutrophil or eosinophil involvement, such as asthma. Our findings suggest that linker histones represent another layer of complexity to the fascinating process of immune cell differentiation.

## Author contributions

GS and AZ designed the study and wrote the manuscript. GS designed and performed experiments and analyzed data, RS provided conceptual input, analyzed and visualized RNA-seq data, BC analyzed the CRISPR/Cas9 screen, AIS provided H1.2/H1.4-deficient mice. All authors commented on the manuscript.

## Acknowledgements

We thank Dr. Volker Brinkmann and Christian Goosmann for their help with electron microscopy and we thank the Max Planck Genome-centre Cologne for performing the sequencing of the PLB-985 samples in this study. We also thank Dr. Alf Herzig, Dr. Borko Amulic and Anna Zychlinsky Scharff for reading and commenting on the manuscript. GS was funded by an Early Postdoc.Mobility and an Advanced Postdoc.Mobility fellowship from the Swiss National Science Foundation. The project was funded by the Max Planck Society.

## Data availability

All relevant data are included in the figures and the supplementary material, RNA-seq raw data are available upon request.

---

**Supplementary Fig. 1.**
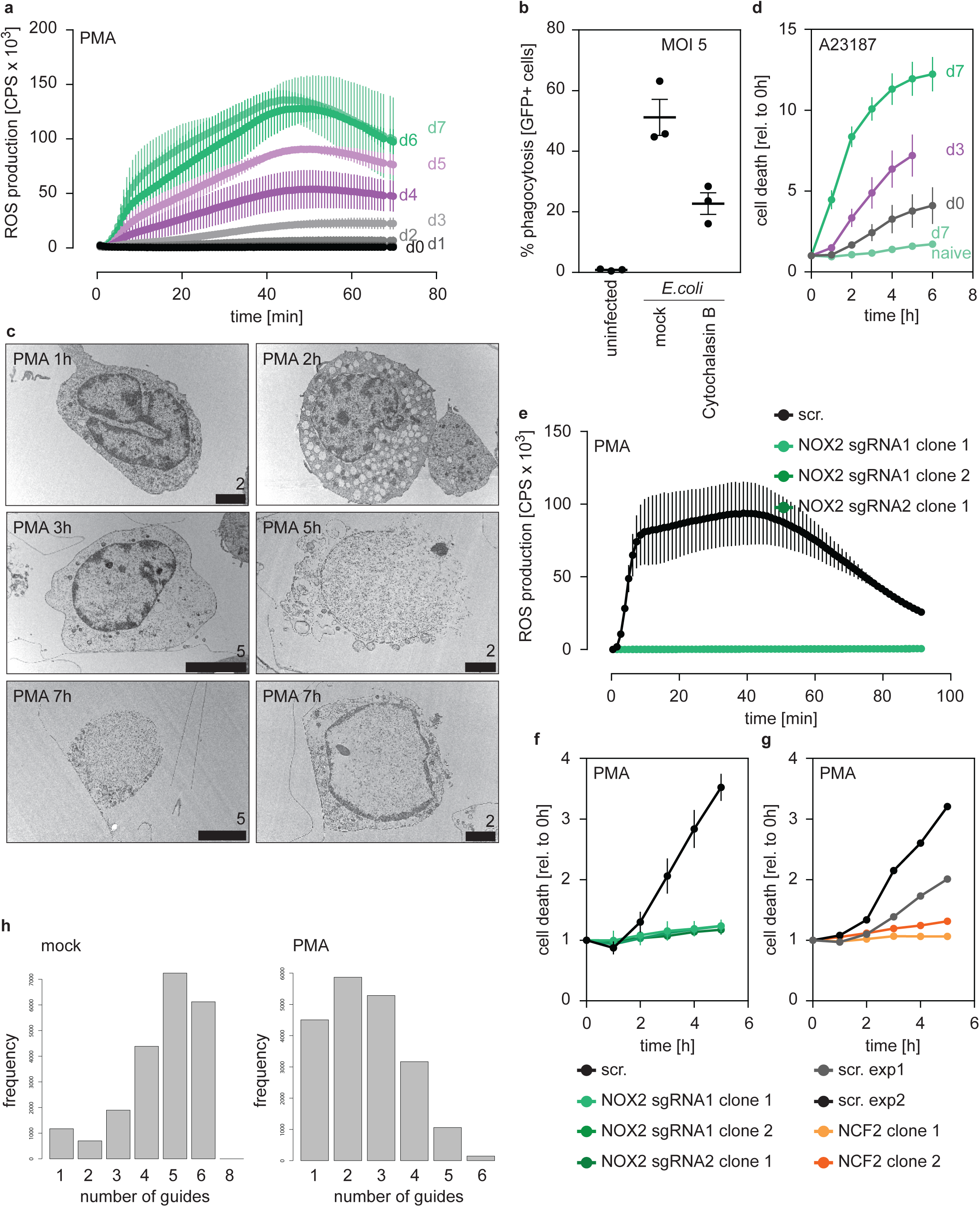
Characterization of PLB-985 function and morphology. **a** ROS production of PLB-985 in response to PMA at different time points of differentiation, depicted are mean -/+ SEM of two independent experiments. **b** Phagocytosis assay of differentiated (d7) PLB-985 incubated with GFP-expressing *E. coli* (multiplicity of infection (MOI): 5), phagocytosis was analyzed by flow cytometry of GFP-positive cells. The phagocytosis inhibitor cytochalasin B (Sigma, 5 μM) was used as a control. **c** TEM images of differentiated PLB-985 (d7) stimulated with PMA for the indicated time points, demonstrating nuclear expansion and, in some cases (5h example), nuclear rupture and chromatin release. Scale bars correspond to indicated values in μm. **d** Cell death in response to the calcium ionophore A23187 (5 μM) at different time points of differentiation, measured by SYTOX Green fluorescence. Depicted are mean -/+ SEM of 3 independent experiments, “d7 naive” are differentiated, but untreated cells and are the same values as in Figure 1D. **e** ROS production of independent NOX2 (*CYBB*) -/- clones (derived from 2 sgRNAs) after treatment with PMA. **f** Cell death of independent NOX2 -/- clones (derived from 2 sgRNAs) after treatment with PMA. (**e, f**) depicted is the mean -/+ SEM of 2-3 independent experiments. (**g**) Reduced cell death of two clones deficient in NCF2 in response to PMA, depicted is the mean of 1 experiment per clone and the scrambled (scr.) controls for the respective experiments. (**h**) sgRNA distribution of PLB-985 after library transduction and differentiation. The left panel shows all sgRNAs identified in the mock-treated sample and demonstrates that mostly 4-6 sgRNAs per gene were identified, indicating good library coverage. The right panel shows the PMA-treated condition where fewer sgRNAs were identified and a left shift was observed, demonstrating selection pressure.

**Supplementary Fig. 2.**
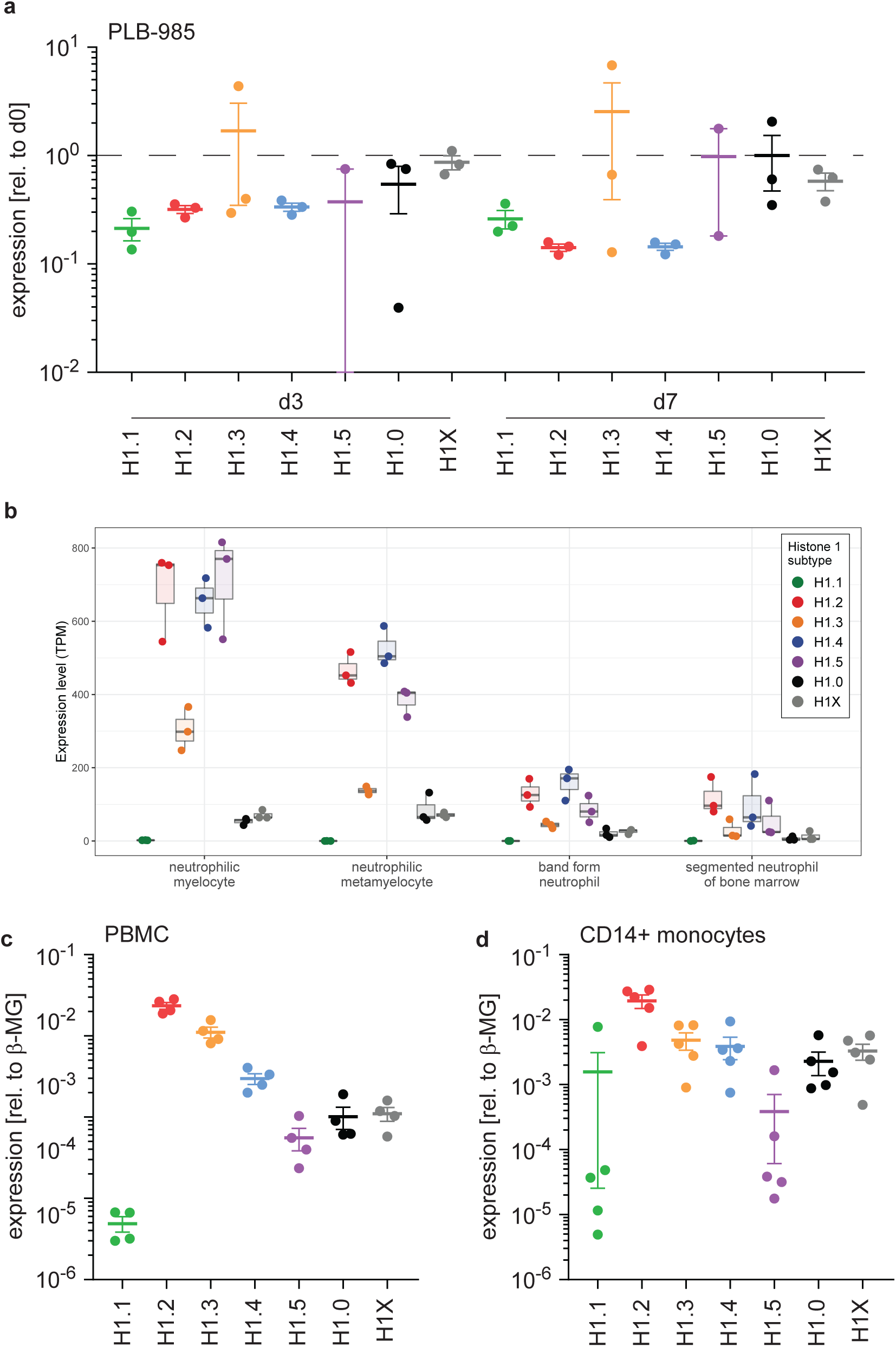
H1 expression levels in human PBMCs, monocytes and PLB-985. **a** Relative mRNA expression levels corresponding to Fig. 2a, values are relative to d0 of differentiation and indicate reduction of mRNA expression for H1.1, H1.2, H1.4 and H1.5 during differentiation. Dashed line indicates unchanged expression. Note that the distribution of samples varied much more in the lowest expressed subtypes H1.3 and H1.5. **b** Transcripts per million (TPM) for the indicated H1 subtypes were calculated for human primary bone marrow cells^27^ at indicated (increasing from left to right) stages of differentiation. mRNA for somatic H1 subtype decreases with differentiation, reflecting exit of cell cycle. **c, d** mRNA expression levels of indicated H1 subtypes in human primary peripheral blood mononuclear cells (PBMC) (**c**) or in MACS-sorted CD14+ human primary monocytes (**d**). Each dot represents a healthy donor, error bars are -/+ SEM.

**Supplementary Fig. 3.**
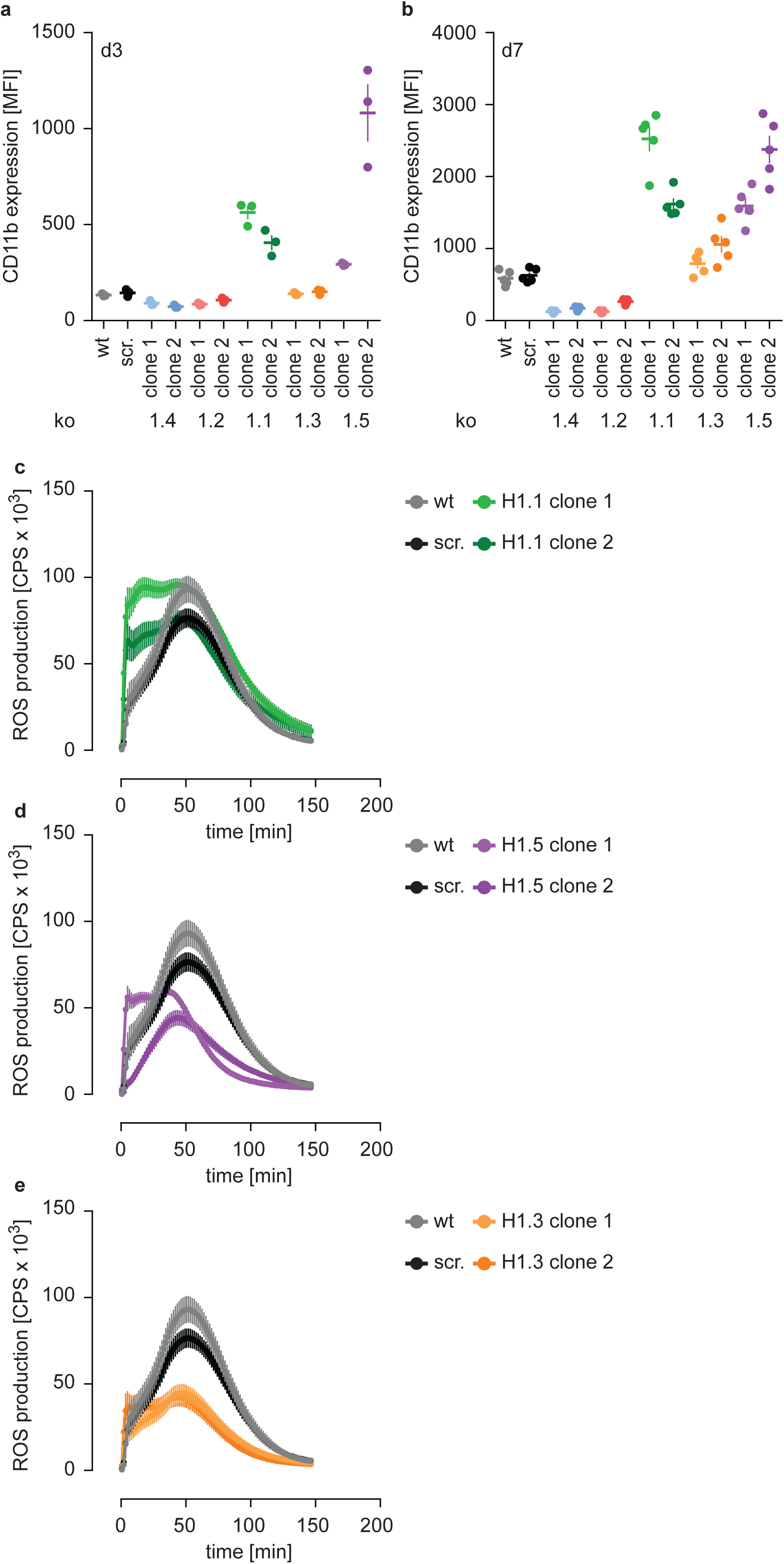
Subtype-specific H1 impact on PLB-985 maturation and function. **a, b** Flow cytometry analysis corresponding to Fig. 3a, b, depicted is the mean fluorescence intensity (MFI) of CD11b in indicated samples at d3 (**a**) or d7 (**b**) of differentiation. Each dot represents an independent experiment; error bars are -/+ SEM. **c-e** ROS production of indicated clones in response to PMA. Depicted is the mean -/+ SEM of 5 independent experiments. wt and scr. values are the same as in Fig. 2d.

**Supplementary Fig. 4.**
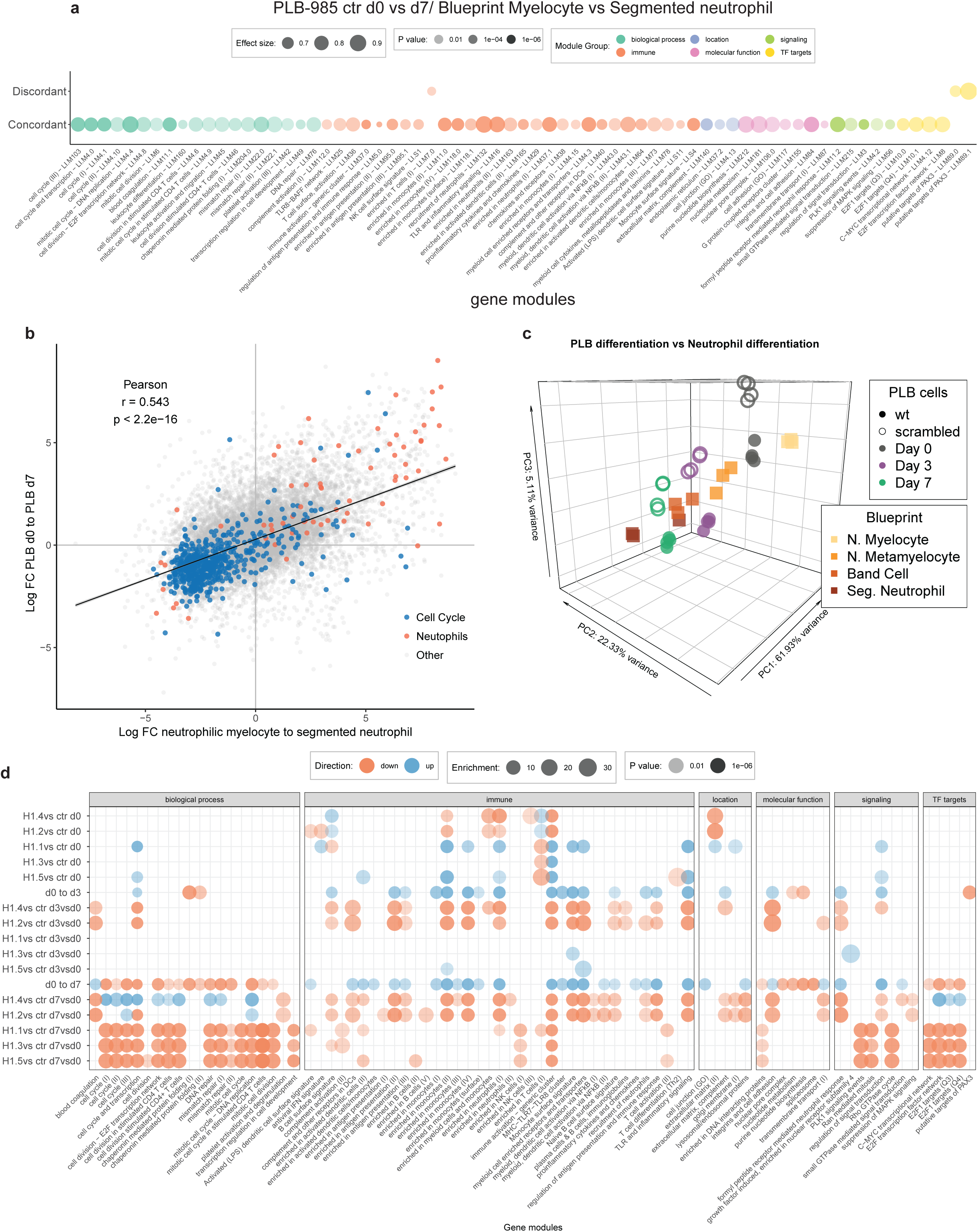
Transcriptional differences between H1-deficient clones during differentiation. **a** Concordantly and discordantly regulated modules when comparing PLB-985 differentitation (wt and scr., d0 to d7) and human neutrophil differentiation (Blueprint, neutrophilic myelocyte to segmented neutrophil of bone marrow) using discordance scores and tmod. Same values as in Fig. 4a, showing the annotated names of modules. **b** Scatter dot plot showing all genes (each dot represents one gene, colors refer to genes belonging to the aggregated neutrophil or cell cycle modules) and their regulation in differentiating PLB-985 compared to human primary cells, demonstrating that neutrophil-annotated modules and cell cycle-annotated modules are mostly regulated in a concordant manner. **c** Principal component analysis of control (wt and scr.) samples compared to human primary RNA-seq data of bone marrow neutrophil precursors (Myelocytes, Metamyelocytes, Band cells and segmented neutrophils of the bone marrow), derived from the blueprint consortium, showing a similar developmental trajectory of primary cells and PLB-985. **d** Gene module enrichment analysis of indicated genotypes; for mutant differentiation, the relative changes of gene module enrichment when compared to control cell differentiation is depicted.

**Supplementary Fig. 5.**
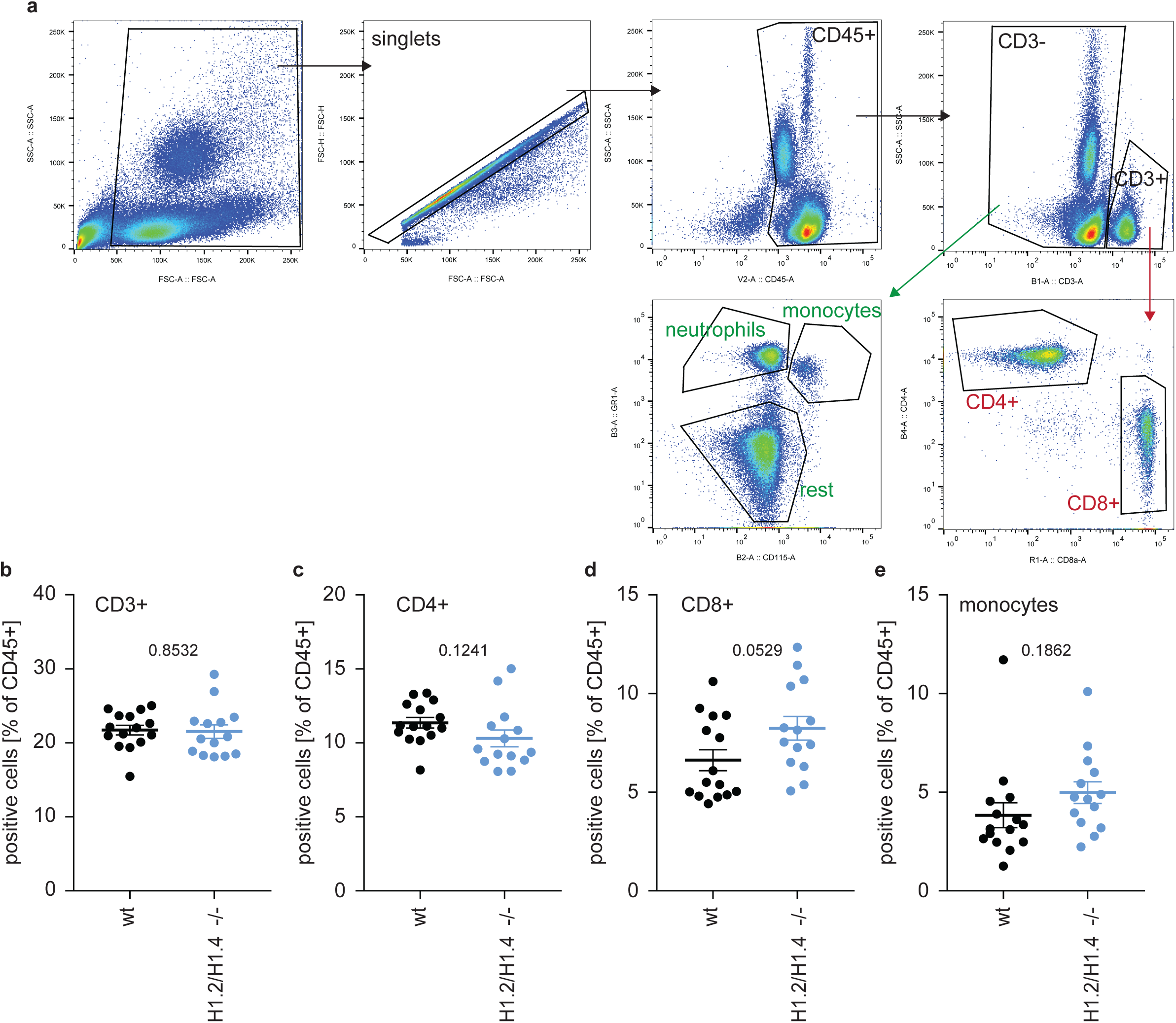
Normal circulating leukocytes in adult H1.2/H1.4 double-deficient mice. **a** Gating strategy for different subsets of immune cells in whole blood. Single cells are gated for expression of CD45, then for CD3. CD3 positive cells are divided into CD4 and CD8 positive T cells, CD3 negative fractions are divided into neutrophils (GR1 positive, CD115 negative), monocytes (GR1 positive CD115 positive) and “rest” (B cells, NK cells and others). **b-e** Abundance of the indicated leukocytes in whole blood of wild type (wt, n=15) and H1.2/H1.4 double-deficient (H1.2/H1.4 -/-, n=14) animals, each dot represents a mouse, error bars are mean -/+ SEM. p values are indicated and derived from an unpaired, two-tailed t test. The profile of circulating immune cells in H1.2/H1.4 -/- animals looks normal.

**Supplementary Fig. 6.**
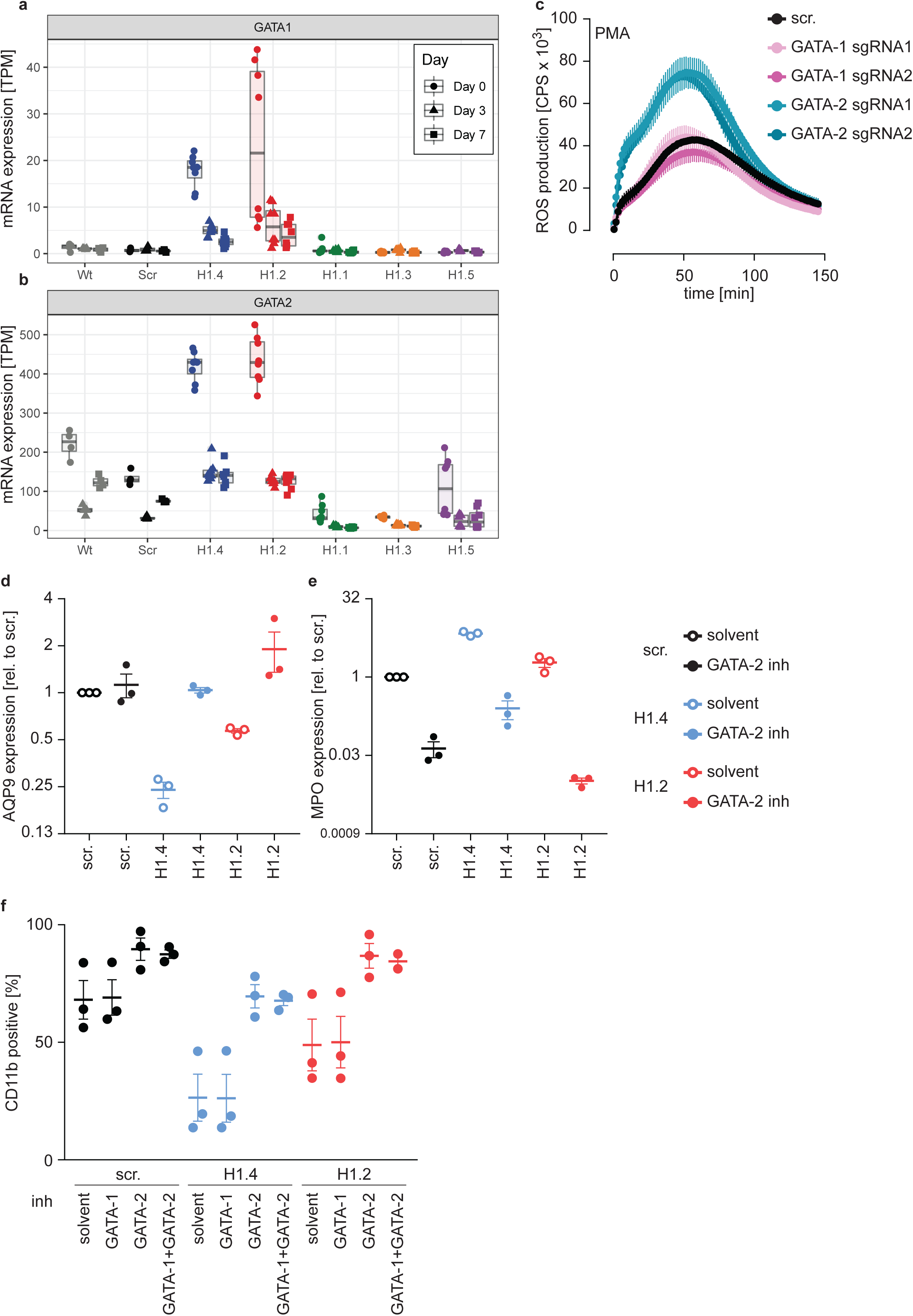
H1.2 or H1.4 deficiency can be reverted by inhibition of GATA-2. **a, b** Transcripts per million (TPM) were calculated from RNA-seq data for GATA-1 (**a**) and GATA-2 (**b**) for the indicated genotypes at all time points of differentiation. GATA-1 expression is enhanced in H1.2 and H1.4 knockout lines at the onset of differentiation. GATA-2 expression is generally higher than GATA-1 and further elevated in knockout lines for H1.2 and H1.4. **c** PLB-985 were transduced with 2 sgRNAs against GATA-1 and 2 sgRNAs against GATA-2. Cells were differentiated in batches and ROS production in response to PMA was measured at d4 of differentiation. Shown is the mean -/+ SEM of 3 independent experiments. **d, e** Treatment of PLB-985 cells with GATA-2 inhibitor during differentiation improves maturation of cells as shown by increased mRNA levels of AQP9 (**d**) and decreased mRNA levels of MPO (**e**). The effect was particularly strong in knockout lines for H1.2 and H1.4. Shown is the mean -/+ SEM of 3 independent experiments. **f** CD11b expression of PLB-985 of the indicated genotypes at d7 of differentiation. Cells were treated with solvent or the indicated inhibitors before differentiation and at d4. Same experiments as in Fig. 6e, including treatment with a combination of inhibitors. Shown is mean -/+ SEM of 3 independent experiments.

**Supplementary Table 1. List of genes identified as hits in the CRISPR/Cas9 screen**

The table shows gene ID and the fold representation of identified sgRNAs in PMA-treated versus untreated PLB-985. sgRNAS that were not identified in either sample are labelled as NA. The table also shows the median and mean values per gene and the percentage of overrepresented (at least 2 fold) sgRNAs.

**Supplementary Table 2. RNA-seq expression tables**

The individual sheets display expression values of indicated conditions (Cont. is wild type and scr. combined, for the respective H1 subtypes values of 2 clones are combined).

**Supplementary Table 3. List of sgRNA sequences, primer sequences and antibodies**

Individual sheets contain sequences of sgRNAs, sequencing primers, qRT-PCR primers and antibody catalog and lot numbers.

